# Lentivirus Transduced T Cell Lines Response to Pro-inflammatory and Antiviral Cytokines in the Presence of HIV

**DOI:** 10.1101/2024.07.27.605446

**Authors:** Kevin Ventocilla, Daniel Miranda, David Jesse Sanchez

## Abstract

Various studies have attempted to understand HIV infection under a diverse range of stimulants including cytokine stimulation. Pro-inflammatory cytokines, such as TNF-α, have been shown to reactivate HIV latency by inducing NF-κB mediated activation of the HIV LTR (long terminal repeats) that contain κB transcriptional binding sites. Interferon-alpha (IFN-α), an anti-viral cytokine, is not well studied as an inducer of HIV activation. However, previous work from our group has shown that HIV can block IFN-α signaling in CD4+ T cells presumably to allow for further viral replication. Initially using HEK 293T cells, we moved to CD4+ T cells lines to develop a system to determine how stimulation with different cytokines impacts signaling within T cell lines. We confirmed that in our system TNF-α triggers activation of NF-κB driven reporters but not in the presence of HIV. In addition, we show that the presence of HIV blocks IFN-α signaling. Taken together, our system demonstrates that HIV by TNF-α, will continue to block IFN-α signaling preventing it from impacting HIV activation. This system can now be used to screen for cytokine based and other molecule activators that may be influenced by the presence of HIV.

## Introduction

The progression of HIV infection in a person living with HIV is dependent the balance of viral replication and the immune response. However, people living with HIV infection, even if they are on antiretroviral therapy (ART) may still develop latent infections of replication-competent virus that are silent amongst resting cells causing HIV reservoirs. Viral latency of HIV is primarily established in activated CD4 T cells as they begin to transition into their long-lived resting state. Infected cells are not actively producing viruses; thus, they are unable to be detected by immune surveillance which relies on the presentation of viral antigen. In addition, as ART targets active viral enzymes, ART will not affect these latently infected cells. Making things more challenging is that there are no known cellular biomarkers that can distinctively separate latently infected cells from uninfected cells. This dilemma makes reducing HIV reservoirs a challenging obstacle for HIV eradication [1].

The formation of HIV latency depends on various events that can be categorized into two groups. One group inhibits the formation of the viral reservoir while the second group promotes the formation of the HIV reservoir. In terms of the first group inhibiting the formation of the viral reservoir, a key regulatory molecule controlling lymphoid tissue homeostasis, B-cell lymphoma 2 (BCL2) also affects HIV homeostasis in infected lymphocytes. BCL2 can induce apoptosis by binding to a procaspase fragment thus affecting the viral reservoir by induction of apoptosis [2]. Another aspect which inhibits the formation of latent HIV reservoir is the presence of the HIV Tat protein which can inhibit the establishment of latent HIV. Tat is an HIV protein that is a potent activator of viral transcription, however it can combat mechanisms for establishing HIV latency by inducing nuclear translocation of active NF-κB thereby keeping HIV transcription active [3].

The second group determining HIV latency has the potential to promote the formation of the viral reservoir. One method of this is through IFN-gamma-induced protein (IP) – 10 to C-X-C chemokine receptor 3 (CXCR3). IP-10 stimulation in resting helper T cells triggers both actin dynamics along with cofilin activation which promotes HIV entering latency [4]. The increase of cofilin and activities by actin promote nuclear migration in resting T cells as well as viral entry.

HIV latency can be reactivated through various ways including cytokine stimulation. Cytokines can also play a vital role in exacerbating the formation of HIV latent virus reservoirs. Pro-inflammatory cytokines can promote viral persistence during ART by upregulating HIV replication [5]. The pro-inflammatory cytokine, TNF-α, has been shown to reactive HIV latency through the NF-κB pathway [6]. This activation is caused by induction of NF-κB to translocate into the nucleus and bind to κB sites in the LTR [7]. However, the antiviral cytokine, IFN-α, and the potential effects it has on HIV latency reversal has not been well studied. Our studies have found that HIV blocks both production of IFN-α and IFN-α signaling in CD4+ T cells [8; 9]. Pursuing latency reactivation triggers such as cytokines including IFN-α could help us better understand what cytokines induce latency reactivation. Here we develop an in vitro system to screen for the activation of reporters by cytokine stimulation while in the presence of the HIV genome.

## Materials and Methods

### Cell Culture and Maintenance

Human Embryonic Kidney 293T cells (HEK 293T) were acquired from ATCC. The HEK 293T cells were maintained with Dulbecco’s modified Eagle medium (DMEM) supplemented with 5% Fetal Bovine Serum and 1% Penicillin-streptomycin. DMEM, Fetal Bovine Serum and Penicillin-Streptomycin was purchased from ThermoFisher Scientific. Cells were incubated in a humidified incubator kept at 37 degrees Celsius with an internal environment supplemented with 5% CO_2_. HEK 293T cells were regularly monitored for growth and sterile conditions.

The Human T Lymphocyte cell line, CCRF-CEM, was purchased from ATCC. The CEM cells were cultured with Roswell Park Memorial Institute (RPMI) medium supplemented with 5% Fetal Bovine Serum and 1% Penicillin-streptomycin. RPMI media was purchased from ThermoFisher Scientific. Cells were incubated in a humid environment at 37 degrees Celsius with an internal environment supplemented with 5% CO_2_. CEM cells were the main cell line used to test transductions and were regularly monitored for growth and sterile conditions.

### HEK 293T Cell Transfections

Transfection was used to measure GFP fluorescence and HIV plasmid expression. HEK 293T cells were transfected in a 24-well plate at a concentration of 0.5 μg of DNA per well. Cells were transfected with the X-tremgene HP DNA transfection reagent (SigmaAldrich) according to the manufacturer’s recommendations. HEK 293T cells were added to a 24-well plate at a concentration of 2.0 x 10^6^ cells/plate. Transfected cells were incubated for 24 hours.

### Luciferase Reporter Lentivirus Assay Protocol

An ISRE or NF-κB Luciferase Reporter Lentivirus (BPS Bioscience) was used to transduce CEM cells to monitor their response towards cytokines. The ISRE lentivirus is a replication incompetent virus that contains a firefly luciferase gene driven by ISRE response element located upstream of the firefly luciferase gene. The use of this virus is intended to monitor the Type I interferon-induced JAK/STAT signaling pathway by measuring luciferase activity. The NF-κB lentivirus is a replication incompetent virus that contains a firefly luciferase gene driven by a NF-κB response element located upstream of the promoter region. The design of this virus is intended to monitor the activation of the NF-kB signaling pathway in target cells by measuring luciferase activity.

To transduce the CEM cell lines, cells were harvested and resuspended in fresh growth media (RPMI containing 5% Fetal Bovine Serum and 1% Penicillin-streptomycin). The cells were diluted to a concentration of 5 x 10^5^ cells/ml in growth media with a final volume of 10 ml. 800 µl of CEM cells were combined with 200 µl of the provided Luciferase Reporter Lentivirus for a final volume of 1 ml into a 1.5 ml microcentrifuge tube. Polybrene was added into the cell-lentivirus mixture at a concentration of 8 µg/ml with a final volume of 0.8 ml. The mixture was gently stirred and incubated at room temperature for 20 minutes. After incubation, the cell-lentivirus mixture was centrifuged for 30 minutes at a speed of 800 x *g* at 32 degrees Celsius. After centrifugation, the media was aspirated and the transduced CEM cell population was resuspended in 2 ml of fresh RPMI growth media. The CEM cells were then left to incubate for 48 hours in humidified incubator at 37 degrees Celsius.

### Cytokine Titration for optimal concentration activation of ISRE firefly luciferase

In this study we will assess ISRE or NF-κB firefly luciferase activity in different cell populations by stimulating cells with the cytokine. Activation of ISRE firefly luciferase is used as an indicator of activation of an antiviral response by IFN stimulation. A titration was performed to determine the optimal concentration of IFN-α needed to induce ISRE firefly luciferase activity in the different cells. We started with an initial stock containing 100,000 Units of IFN-α in this study; units of IFN-α are defined as a measurement of activity for IFN-α which is to modulate immune responses. A titration was established using HEK 293T cell populations with varying concentrations of IFN-α. The baseline concentration of IFN-α is 100 U/µl. Increasing the baseline concentration by 10-fold, the next IFN-α concentration is 1000 U/µl. Our control for this titration will be labeled as Vehicle consisting of an HEK cell population stimulated with vehicle. After a 24-hour stimulation, there was significant difference in ISRE firefly luciferase activity when stimulated with either 100 U/µl or 1000 U/µl IFN-α. After analyzing the minor differences between 100 U/µl or 1000 U/µl concentration of IFN-α, we began to precede the study with 100 U/µl IFN-α concentration.

A titration was performed to indicate the correct concentration of TNF-α needed to induce optimal activation of NF-κB firefly luciferase activity in the cell population. Previous studies have shown that 10 ng/µl is needed to induce an inflammatory cytokine response in a cell population by TNF-α. For this study we have established 10 ng/µl as our baseline concentration for TNF-α. A titration was established using HEK 293T cell populations with varying concentrations of TNF-α. The baseline concentration of TNF-α is 10 ng/µl. Increasing the baseline concentration by 10-fold, the next TNF-α concentration is 100 ng/µl. Our control for this titration will be labeled as 0 ng/µl consisting of an HEK cell population stimulated with vehicle. After 24-hour stimulation, significant difference was found in NF-κB firefly luciferase activity in the HEK cell populations stimulated with either 10 ng/µl or 100 ng/µl TNF-α. We preceded with the study using 100 ng/µl TNF-α concentration to stimulate our cell populations for 24 hours due to the higher luciferase response that is seen in the titration. It is recommended that titrations be done for particular vendors to avoid production differences.

### Cytokine Stimulation

After HEK 293T or CEM cells were successfully transfected with plasmids they were subjected to cytokine stimulation from the anti-viral cytokine, IFN-α and pro-inflammatory cytokine, TNF-α. The full-length HIV plasmids has been engineered to have the GFP ORF driven by the HIV LTR. Cell transfected with the HIV plasmid will be stimulated with cytokines and GFP activity was monitored. Along with the HIV plasmids, cells are co-transfected with Renilla luciferase plasmid that functions as a monitor of cell viability due to the constitutive expression of Renilla. Renilla luciferase levels function as a normalization factor towards the experimental reporter luciferase.

## Results

### HEK 293T cells as in vitro models of cytokine response in the presence of HIV

To see the effects of cytokines in a cell population with HIV gene expression, we used HEK 293T cells due to their robust transfection efficiency. We developed the reporter system consisting of HEK 293T cells transfected with two different HIV plasmids that express GFP under the control of the HIV LTR: (1) NL4-3 HIV plasmid and (2) ΔVpu / ΔNef NL4-3 HIV plasmid. These versions of HIV do not express Env and are thus able to be used to screen without release of infectious HIV. The ΔVpu / ΔNef NL4-3 HIV plasmid does not express either the Vpu or Nef accessory proteins [10].

The HEK 293T cells were stimulated with the cytokines, IFN-α or TNF-α. The TNF-α cytokine is added to induce NF-κB. Of note, HIV LTRs have κB promoter elements and thus should upregulate gene expression of both the NL4-3 and ΔVpu / ΔNef NL4-3 in response to TNF-α. The IFN-α cytokine should induce an upregulation of the antiviral ISRE promotor element in cells. HEK 293T cells that were not treated with the HIV plasmids (labeled as vector) displayed an increase in ISRE firefly luciferase when stimulated with IFN-α (Fig. 1A (gray bars) and 1B). HEK 293T cells transfected with the NL4-3 HIV plasmid had a general decrease in ISRE firefly luciferase when stimulated with IFN-α when compared to the HEK 293T vehicle containing NL4-3 HIV plasmid that were not stimulated with IFN-α cytokine (Fig. 1A and 1C). HEK 293T cells transfected with the ΔVpu / ΔNef NL4-3 HIV plasmid still show a general reduction in ISRE Firefly Luciferase but when stimulated with IFN-α showed an increase in ISRE Firefly Luciferase compared to the HEK vehicle containing the ΔVpu / ΔNef NL4-3 HIV plasmid (Fig. 1A and 1D). This suggests that the accessory proteins Vpu, Nef or both block IFN-α signaling leading to a decrease in ISRE expression, as our group has previously shown.

**Figure 1:**
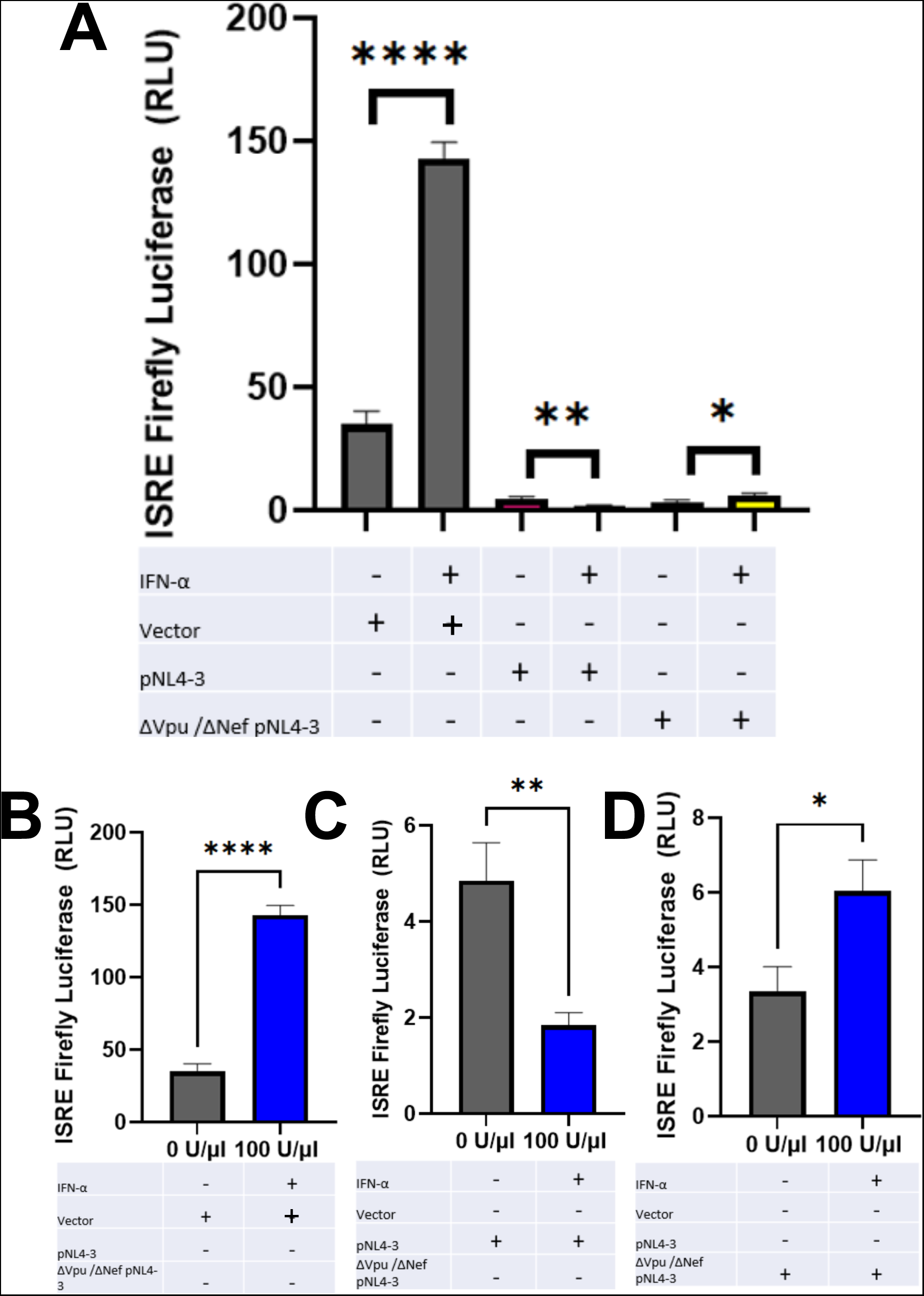
HEK 293T cells transfected with NL4-3 and ΔVpu/ΔNef HIV plasmid stimulated with IFN-α: **(A)** HEK 293T cell population transfected with NL4-3 and ΔVpu/ΔNef NL4-3 HIV plasmid and stimulated with IFN-α. ISRE firefly luciferase is measured by a luminometer. **(B)** HEK vehicle stimulated with 0 U/µl and 100 U/µl IFN-α for 24 – hours. **(C)** HEK cells transfected with NL4-3 HIV plasmid and stimulated with 0 U/µl and 100 U/µl IFN-α for 24 – hours. **(D)** HEK cells transfected with ΔVpu/ΔNef NL4-3 HIV plasmid and stimulated with 0 U/µl and 100 U/µl IFN-α for 24 – hours. Experiments were done in triplicate and ISRE Firefly levels were normalized to Renilla Luciferase. Statistics were determined using unpaired T-test with * being a p value of 0.0239, ** being a p value of 0.0025 and **** being a p value of < 0.0001.

The HEK 293T cell reporter system that we developed was analyzed when stimulated with TNF-α. Vector HEK cells were stimulated with TNF-α they showed a significant increase in NF-κB Firefly Luciferase (Fig. 2A and 2B). HEK cells that were transfected with NL4-3 plasmid were stimulated with TNF-α which showed general increase in NF-κB Firefly Luciferase compared to the vector HEK cell population (Fig. 2A and 2C). The HEK cell population transfected with ΔVpu / ΔNef NL4-3 HIV plasmid was stimulated with TNF-α also showed a general increase in NF-κB Firefly Luciferase compared to the vector cell population. This suggests that Vpu, Nef or both play a role in upregulating NF-κB promoter activity (Fig. 2A and 2D) The transfected HEK cell population was also analyzed by flow cytometry to assess the levels of GFP production which is linked to HIV genome transcription. As seen in Figure 3 significant amount of GFP production is observed indicating successful transfection and active HIV genome transcription [11].

**Figure 2:**
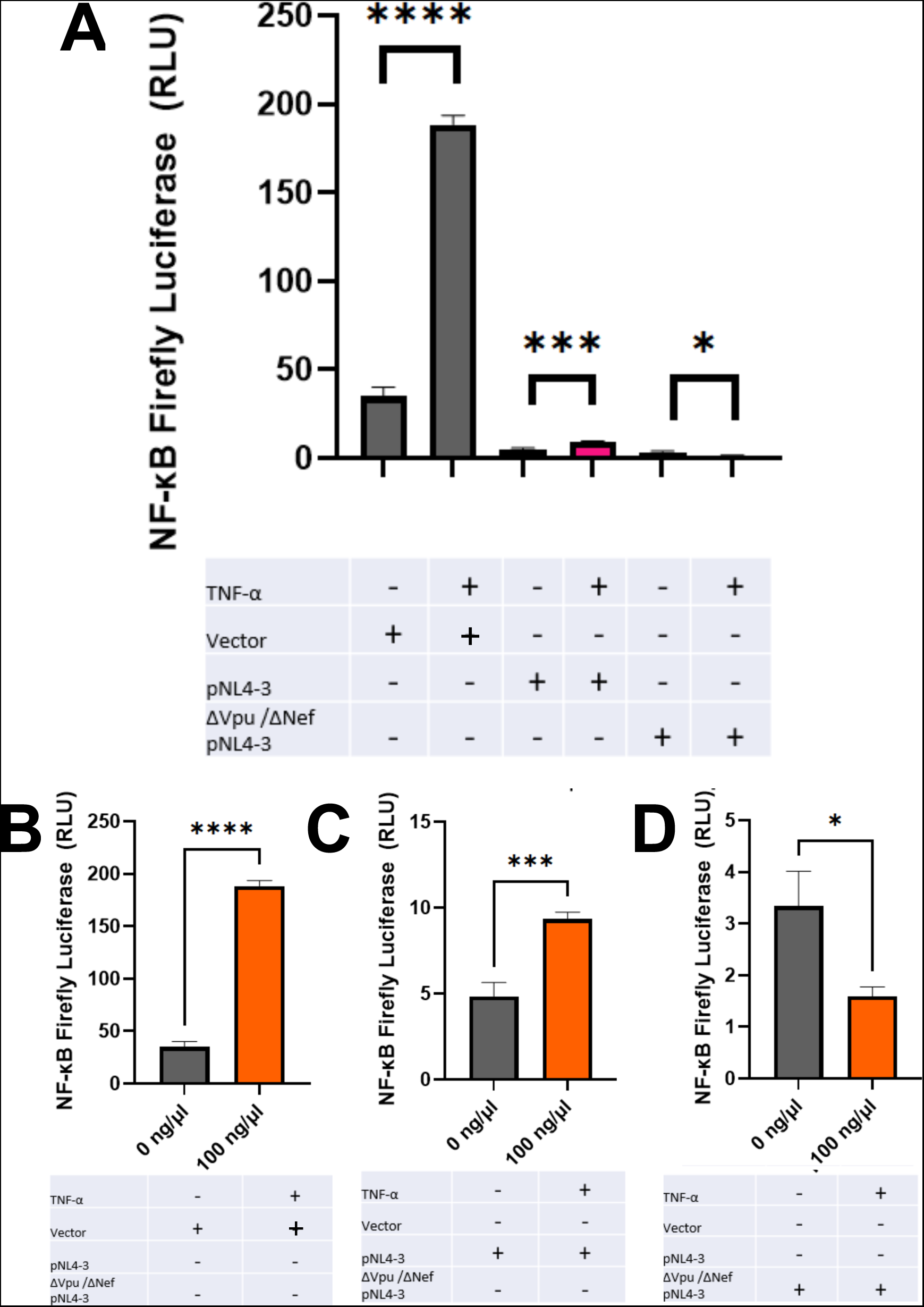
HEK 293T cells transfected with NL4-3 and ΔVpu/ΔNef HIV plasmid stimulated with TNF-α: **(A)** HEK 293T cell population transfected with NL4-3 and ΔVpu/ΔNef NL4-3 HIV plasmid and stimulated with TNF-α. NF-κB firefly luciferase is measured by a luminometer. **(B)** HEK vehicle stimulated with 0 ng/µl and 100 ng/µl TNF-α for 24 – hours. **(C)** HEK cells transfected with NL4-3 HIV plasmid and stimulated with 0 ng/µl and 100 ng/µl TNF-α for 24 – hours. **(D)** HEK cells transfected with ΔVpu/ΔNef NL4-3 HIV plasmid and stimulated with 0 ng/µl and 100 ng/µl TNF-α for 24 – hours. Experiments were done in triplicate and NF-κB Firefly levels were normalized to Renilla Luciferase. Statistics were determined using unpaired T-test with * being a p value of 0.0243, *** being a p value of 0.0001 and **** being a p value of < 0.0001.

**Figure 3:**
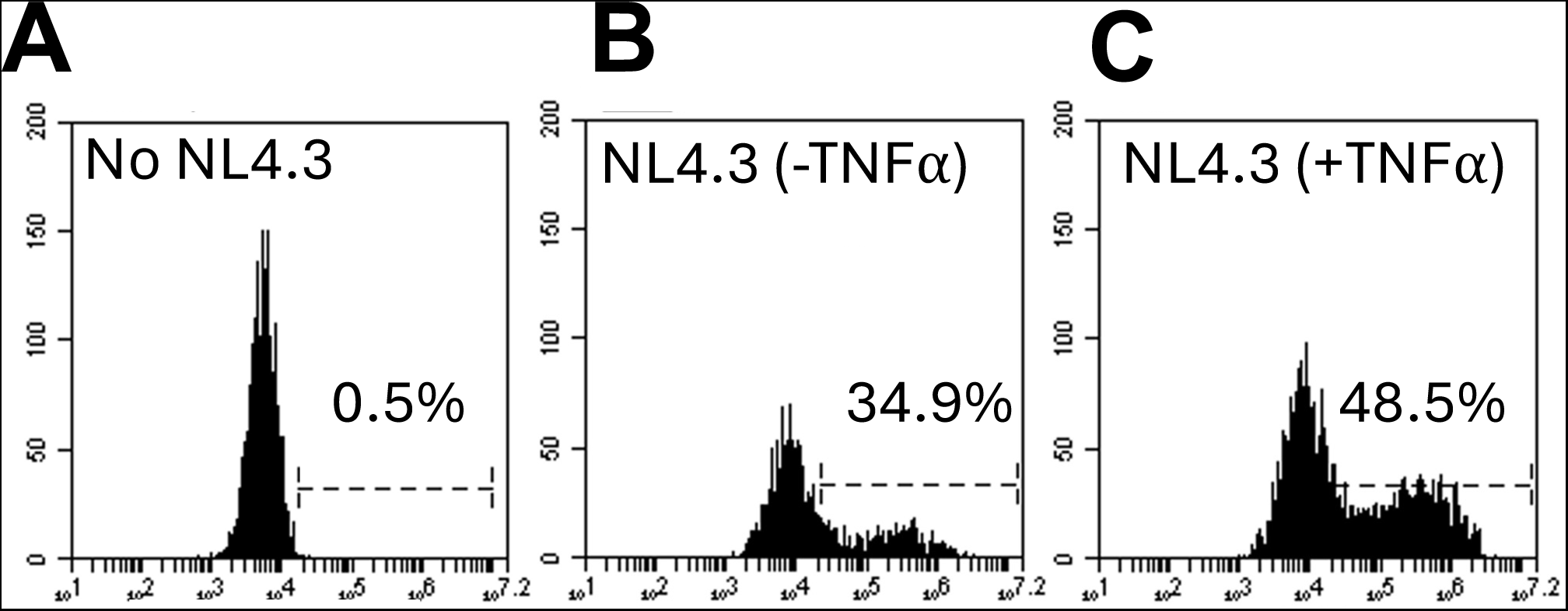
Flow cytometry analysis of HEK 293T cells transfected with NL4-3 Plasmid: **(A)** FL1-A analysis of GFP production from HEK 293T cell vehicle population **(B)** FL1-A analysis of GFP production from HEK 293T cell population transfected with the NL4-3 HIV plasmid **(C))** FL1-A analysis of GFP production from HEK 293T cell population transfected with the NL4-3 HIV plasmid and stimulated with the pro-inflammatory cytokine TNF-α. Experiments were performed in triplicates and analyzed via Flow Cytometry. These are representative images.

### CEM Cells transfection efficiency is not sufficient to monitor cytokine stimulation

Transient transfection of CEM cells was used to assess the effects of cytokines in a T cell population that contained HIV-1 plasmid. CEM cells were transfected with 2 HIV plasmids engineered to express GFP under control of the HIV LTRs: NL4-3 plasmid and ΔVpu / ΔNef NL4-3 HIV plasmid. After transfection, the cell populations were stimulated with IFN-α or TNF-α. Comparing three different CEM cell populations (vector, NL4-3, ΔVpu/ΔNef) after IFN-α stimulation for 24 hrs, no significant difference in ISRE firefly luciferase was found (Fig. 4A). The CEM reporter system was then assessed after TNF-α stimulation for 24 hours. Comparing the three CEM cell populations (vector, NL4-3, ΔVpu/ΔNef) after stimulation, there is no significant difference in NF-kB firefly luciferase (Fig. 4B).

**Figure 4:**
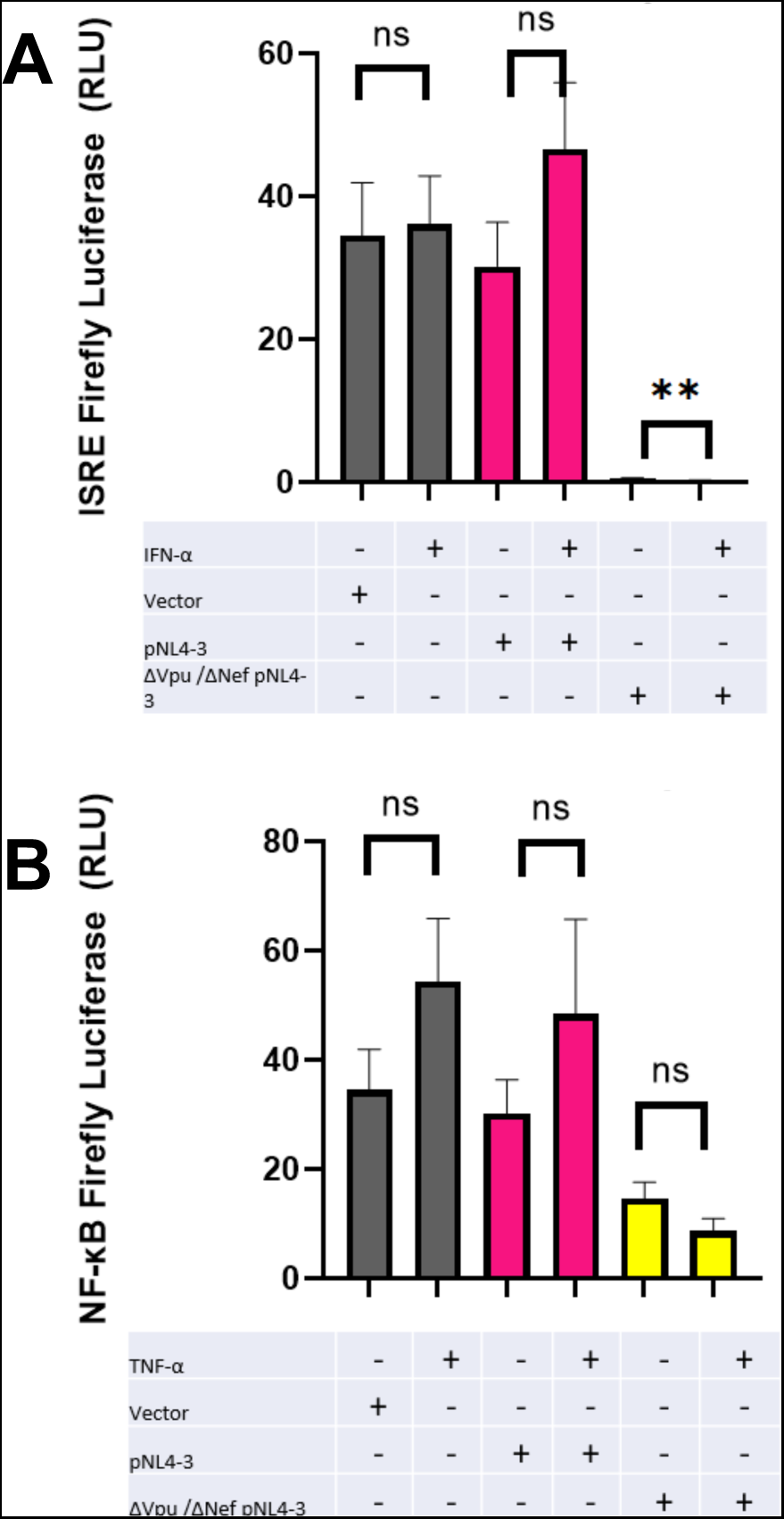
CEM Cells transfected with HIV plasmids and stimulated with Cytokines. CEM cells were transfected with the NL4-3 HIV plasmid and ΔVpu/ΔNef NL4-3 HIV plasmid. Cell populations were stimulated for 24 – hours with **(A)** IFN-α (100 U/µl) or **(B)** TNF-α (100 ng/µl). Experiments were done in triplicate and ISRE Firefly levels were normalized to co-transfected Renilla Luciferase. Statistics were determined using unpaired t-test with ** being a p value of 0.0044

### CEM T cells Transduced with ISRE Luciferase response to IFN-α and TNF-α stimulation

To overcome the limitation of low transfection efficiency in CEM cells, a CEM cell reporter system was constructed with CEM cells that have been stably transduced with an ISRE luciferase reporter lentivirus. The lentivirus contains a firefly luciferase gene that is controlled by an ISRE promoter element allowing monitoring of ISRE inducing signaling in a CEM cell population. We transfected this reporter system to transfected with either a plasmid containing NL4-3 HIV plasmid or the ΔVpu/ ΔNef NL4-3 HIV plasmid. As shown in Figure 5A, CEM cell populations were stimulated with IFN-α for 24 hours were monitored. The ISRE lentivirus-transduced CEM cell population that were not transfected with the HIV plasmids showed significant increase in ISRE firefly luciferase activity when stimulated with IFN-α (Fig. 5A). This shows that the transduced ISRE lentivirus were properly incorporated into the CEM cell population establishing a cell line that is responsive to IFN-α allowing ISRE activation to be monitored by luciferase.

**Figure 5:**
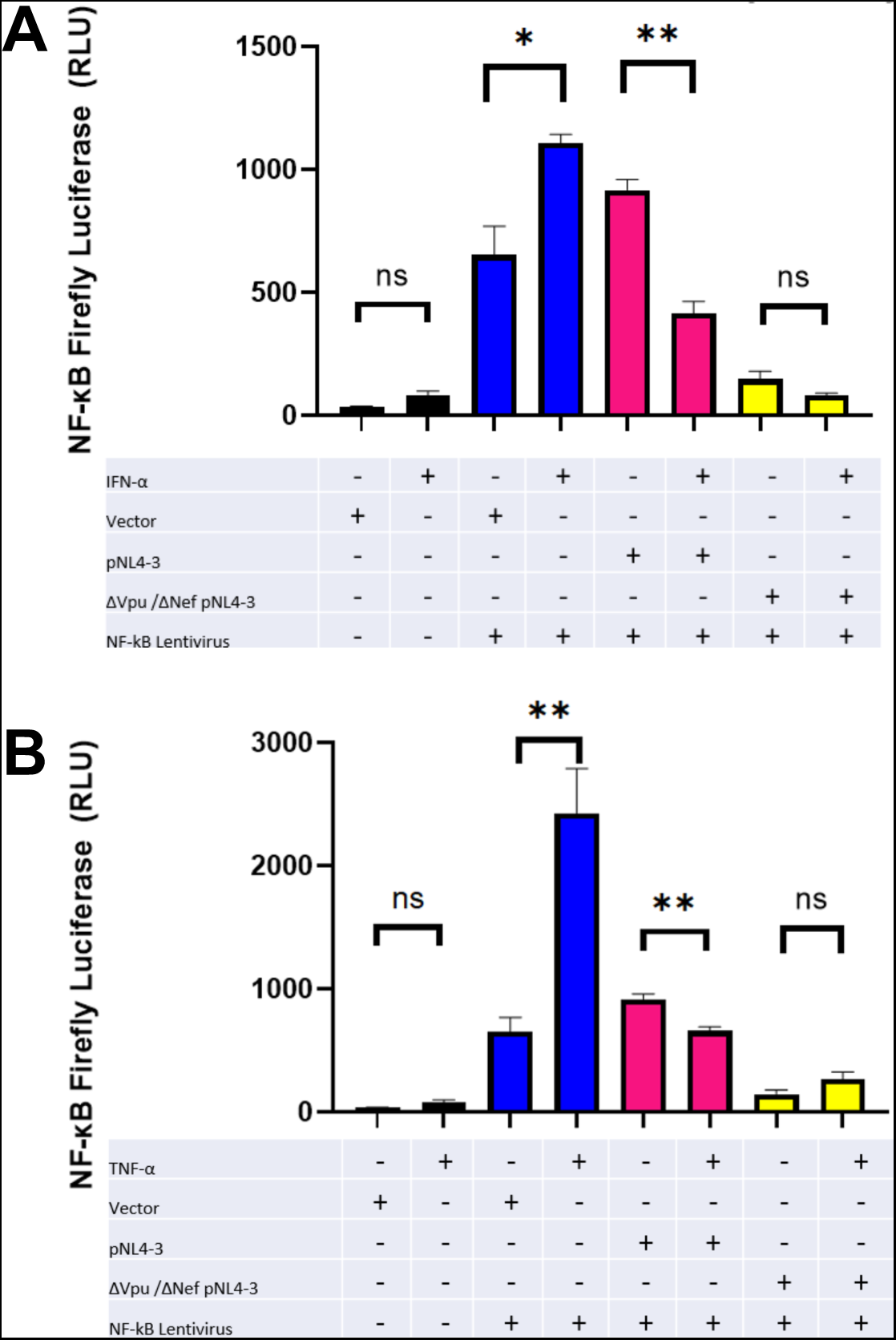
ISRE Luciferase transduced CEM cells stimulated with IFN-α or TNF-α. CEM cells transduced with ISRE Luciferase lentivirus were transfected with either NL4-3 HIV plasmid or ΔVpu/ΔNef NL4-3 plasmid. **(A)** CEM cells were transfected with either NL4-3 HIV plasmid or ΔVpu/ΔNef NL4-3 plasmid. CEM cells were stimulated with IFN-α for 24-hours. Experiments were done in triplicate and ISRE Firefly levels were normalized to co-transfected Renilla Luciferase. Statistics were determined using unpaired t-test with * being a p value of 0.0050 and ** being a p value of 0.020. **(B)** CEM cells were transfected with either NL4-3 HIV plasmid or ΔVpu/ΔNef NL4-3 plasmid. CEM cells were stimulated with TNF-α for 24-hours. Experiments were done in triplicate and ISRE Firefly levels were normalized to co-transfected Renilla Luciferase. Statistics were determined using unpaired t-test with * being a p value of 0.0280 and ** being a p value of 0.0423.

The ISRE lentivirus-transduced CEM population transfected with the NL4-3 HIV plasmid did not display a significant difference in luciferase activity upon stimulated with IFN-α. This suggests that the NL4-3 plasmid inhibits IFN signaling preventing ISRE firefly luciferase activity (Fig. 5A). This is consistent with previous findings from our group that show that HIV inhibits IFN signaling in T cells. However, the ISRE lentivirus-transduced CEM population transfected with ΔVpu/ ΔNef NL4-3 displayed a significant increase in ISRE firefly luciferase activity when stimulated with IFN-α. This shows that transfection with the ΔVpu/ ΔNef NL4-3 HIV plasmid is still able to allow for high amounts of luciferase activity suggesting that the accessory proteins Vpu, Nef, or both play a role in blocking IFN-signaling in the presence of the entire NL4-3 HIV genome (Fig. 5A).

The transduced CEM reporter system was further assessed for ISRE firefly luciferase activity when stimulated with TNF-α for 24 hours. Transduced CEM cells were left as a vehicle control or transfected with the NL4-3 HIV plasmid or ΔVpu/ ΔNef NL4-3 plasmid. After a 24-hour TNF-α stimulation, a significant increase in ISRE firefly luciferase activity was observed in the transduced CEM vehicle population while a significant decrease was seen in cells transfected with the NL4-3 HIV plasmid (Fig. 5B). This suggests that HIV blocks TNF-α signaling that could lead to ISRE activation. The ISRE Luciferase lentivirus transduced CEM population that was transfected with the ΔVpu/ ΔNef NL4-3 plasmid does not show a significant difference in ISRE firefly luciferase activity when stimulated with TNF-α for 24 hours. The increase in overall activation suggests that Vpu and/or Nef may be interacting with this pathway.

### CEM T cells Transduced with κB Luciferase response to IFN-α and TNF-α stimulation

A CEM cell reporter system was constructed with CEM cells that have been transduced with an NF-κB luciferase reporter lentivirus. The lentivirus contains a firefly luciferase gene that is driven by four NF-κB response elements in the promoter region. The purpose of this lentivirus is to monitor TNF-α-induced signaling in a CEM cell population in the presence of HIV. This reporter system consisted of CEM cells transduced with NF-κB luciferase reporter lentivirus that is then transfected with either NL4-3 HIV plasmid or ΔVpu/ ΔNef NL4-3 HIV plasmid.

In Figure 6A, three CEM cell populations were stimulated with IFN-α cytokines for a duration of 24 hours and analyzed by a luminometer. The NF-κB lentivirus that was not transfected with HIV plasmid showed a significant increase in luciferase activity when stimulated with IFN-α. The second CEM cell population to be stimulated with IFN-α was the CEM cells transfected NL4-3 HIV plasmid; significant decrease in luciferase activity was seen within this population. The transduced CEM cell population transfected with ΔVpu/ ΔNef NL4-3 HIV plasmid was stimulated with IFN-α resulting in no significant difference in luciferase activity.

**Figure 6:**
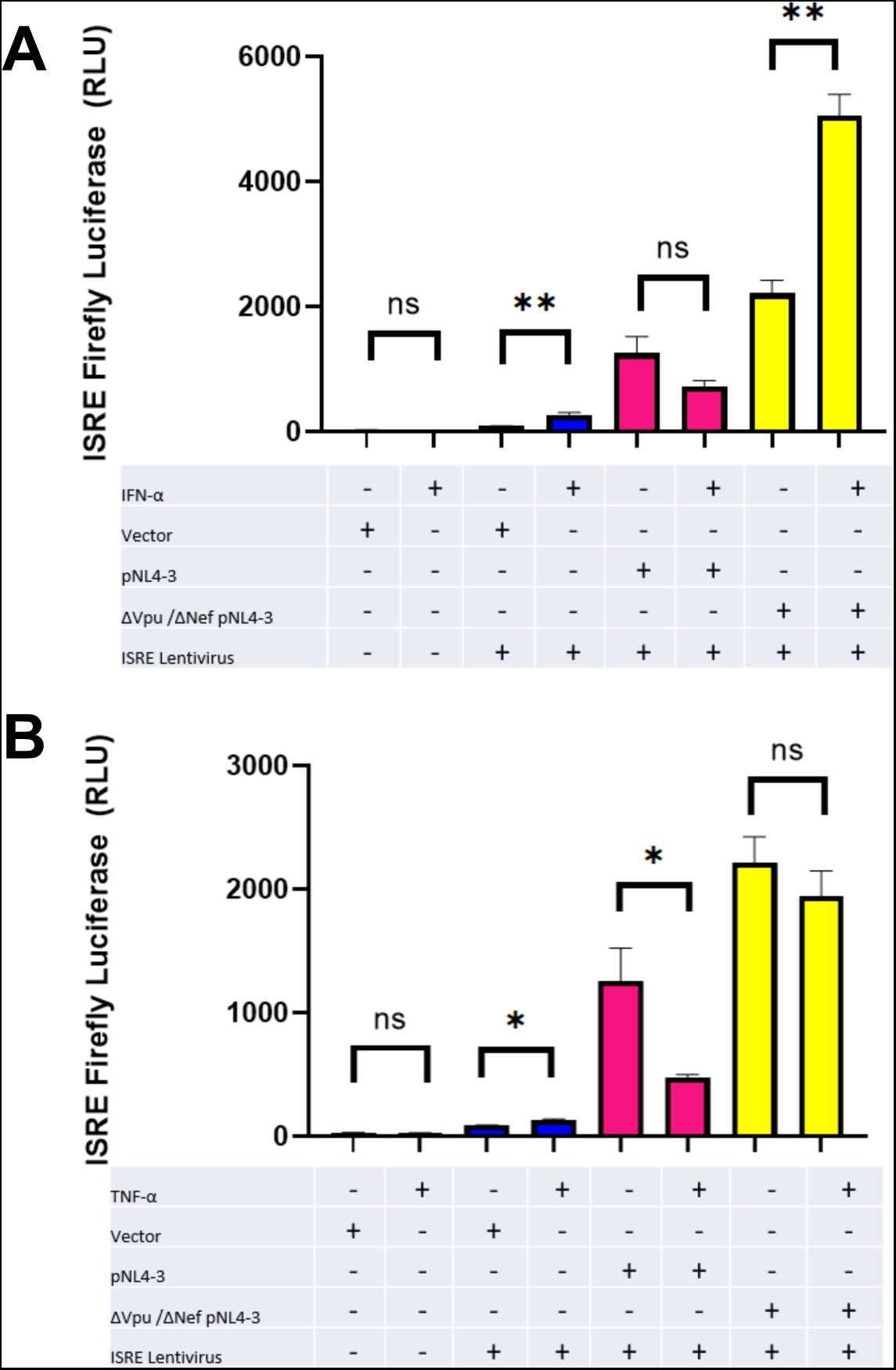
κB Luciferase transduced CEM cells stimulated with IFN-α or TNF-α. CEM cells transduced with κB Luciferase lentivirus were transfected with either NL4-3 HIV plasmid or ΔVpu/ΔNef NL4-3 plasmid. **(A)** CEM cells were stimulated with IFN-α for 24-hours. Experiments were done in triplicate and NF-κB Firefly levels were normalized to co-transfected Renilla Luciferase. Statistics were determined using unpaired T-test with * being a p value of 0.0212 and ** being a p value of 0.0014. **(B)** Cells were stimulated with TNF-α for 24-hours and luciferase was measured. Experiments were done in triplicate and κB Firefly levels were normalized to co-transfected Renilla Luciferase. Statistics were determined using unpaired T-test with ** being a p value ≤ 0.0100.

The CEM reporter system was assessed for NF-κB firefly luciferase activity after a 24-hour stimulation period with TNF-α. The reporter system consists of three different CEM cell populations. The CEM cell population that has been transduced with NF-κB Luciferase Reporter Lentivirus but not transfected with HIV plasmids was stimulated with TNF-α and results were analyzed by a luminometer. The findings showed there was a significant difference in luciferase activity as shown in Figure 6B. The second CEM cell population stimulated with TNF-α for 24 hours was transfected with NL4-3 HIV plasmid; where a slight decrease in luciferase activity was observed and recorded. The third transduced CEM cell population stimulated with TNF-α was transfected with ΔVpu/ ΔNef NL4-3 HIV plasmid and measured luciferase activity finding no significant difference.

## Discussion

The pro-inflammatory cytokine TNF-α has been shown to reactivate dormant HIV-1 provirus in HIV-infected cells to varying degrees. Therefore, TNF-α has been hypothesized as a promising agent for reactivating latent HIV [12]. However, its therapeutic use remains controversial due to its toxic effects at high levels. Our data confirm that TNF-α triggers the activation of HIV transcription in HEK 293T cells transfected with the NL4-3 HIV plasmid. In contrast, no significant difference in activity was observed in CEM cells transfected with the NL4-3 HIV plasmid, possibly due to lower transfection efficiency in CEM cells compared to HEK 293T cells.

To better utilize a T cell line, we performed a transduction of CEM cells with three different lentiviruses: (1) a control empty lentivirus delivering a luciferase reporter without promoter elements, (2) an ISRE luciferase lentivirus with the ISRE promoter element in the HIV LTR driving luciferase, and (3) an NF-κB luciferase lentivirus with the κB promoter element in the HIV LTR driving luciferase. The ISRE luciferase lentivirus allows monitoring of ISRE activation by Type I interferons such as IFN-α.

Upon successful transduction, the CEM cells were stimulated with TNF-α for 24 hours, resulting in a significant increase in ISRE firefly luciferase activity. This finding suggests that TNF-α may indirectly induce ISRE activation, potentially through pathways independent of JAK-STAT signaling, as previously reported. In contrast, CEM cells transfected with the NL4-3 HIV plasmid showed a reduction in ISRE activity upon TNF-α stimulation, while those transfected with the ΔVpu/ΔNef NL4-3 plasmid showed no significant difference in ISRE activity. This indicates that the accessory proteins Vpu and/or Nef may interfere with this pathway.

We also examined the response of ISRE luciferase-transduced CEM cells to IFN-α stimulation. In the absence of HIV plasmids, these cells showed a significant increase in ISRE firefly luciferase activity upon IFN-α stimulation. This response confirms proper integration of the ISRE luciferase lentivirus. However, CEM cells transfected with the NL4-3 HIV plasmid did not display a significant difference in ISRE activity when stimulated with IFN-α, suggesting that HIV inhibits IFN signaling in T cells, consistent with our previous findings.

Interestingly, the ΔVpu/ΔNef NL4-3 transfected CEM cells showed a significant increase in ISRE firefly luciferase activity when stimulated with IFN-α. This highlights the role of Vpu and Nef in blocking IFN signaling, as their absence allows for higher ISRE activity.

For NF-κB luciferase-transduced CEM cells, significant luciferase activity was observed following TNF-α stimulation, supporting the activation of the NF-κB pathway. However, CEM cells transfected with the NL4-3 HIV plasmid exhibited lower NF-κB activity upon TNF-α stimulation, likely due to the inhibitory effects of Vpu on the NF-κB pathway. The ΔVpu/ΔNef transfected CEM cells, however, showed no significant difference in luciferase activity between the vehicle and TNF-α stimulated populations.

In conclusion, our study demonstrates that HIV gene expression is enhanced in a pro-inflammatory microenvironment. Understanding the cytokine-driven reactivation of latent HIV can inform future therapeutic strategies. Notably, our findings emphasize the inhibitory roles of Vpu and Nef on Type I interferon signaling, providing insights into the mechanisms by which HIV evades the host immune response. This system can now be utilized to screen for activators of pro-inflammatory or antiviral responses in the context of HIV, potentially guiding the development of latency reactivation therapeutics.

## ACKNOWLEDGMENT

Research was supported by NIH Grants R15AI138847, R03DE023307 and R21CA261297 to DJS. The granting agency had no role in overall study design, the design, analysis, and interpretation of the data, decision to publish, or preparation of the manuscript.

The following reagent was obtained through the NIH HIV Reagent Program, Division of AIDS, NIAID, NIH: Human Immunodeficiency Virus (HIV-1), Strain NL4-3 3’ ΔVpu ΔNef Partial Molecular Clone (p230-11), ARP-2535, contributed by Dr. Ronald Desrosiers, Dr. Jim Gibbs and Dean Regier. The following reagent was obtained through the NIH HIV Reagent Program, Division of AIDS, NIAID, NIH: Human Immunodeficiency Virus 1 (HIV-1) NL4-3 ΔEnv EGFP Reporter Vector, ARP-11100, contributed by Dr. Haili Zhang, Dr. Yan Zhou and Dr. Robert Siliciano.

## AUTHOR CONTRIBUTIONS

DJS and DMJr were involved in the conception of the project. KV, DMJr, and DJS were involved in the design, analysis, and interpretation of the data. DJS and KV were involved in drafting of the paper. All authors provided final approval of the version to be published; In addition, all authors agree to be accountable for all aspects of the work.

## DATA AVAILABILITY STATEMENT

The data that support the findings of this study are available from the corresponding author, DJS, upon reasonable request.

